# Signalling pathway crosstalk stimulated by L-proline drives differentiation of mouse embryonic stem cells to primitive-ectoderm-like cells

**DOI:** 10.1101/2023.02.14.528585

**Authors:** Hannah J. Glover, Holly Holliday, Rachel A. Shparberg, David Winkler, Margot Day, Michael B. Morris

**Affiliations:** School of Medical Sciences, University of Sydney, Australia; Department of Biochemistry and Chemistry, Latrobe Institute for Molecular Science, Latrobe University, Australia; Monash Institute of Pharmaceutical Sciences, Monash University, Australia; Advanced Materials and Healthcare Technologies, School of Pharmacy, University of Nottingham, UK

**Author notes:** Naomi Berrie Diabetes Center, Columbia Stem Cell Initiative, Department of Pediatrics, Columbia University Irving Medical Center, New York, NY. Children’s Cancer Institute, Lowy Cancer Research Centre, University of New South Wales, Australia (Holly Holliday). School of Clinical Medicine, University of New South Wales, Australia (Holly Holliday).

**Keywords:** L-proline, amino acid, primitive ectoderm, cell signalling, mouse embryonic stem cell, growth factor

## Abstract

The amino acid L-proline exhibits novel growth factor-like properties during development - from improving blastocyst development to driving neurogenesis *in vitro*. Addition of 400 μM L-proline to self-renewal medium drives mouse embryonic stem cells (ESCs) to a transcriptionally distinct pluripotent cell population - early primitive ectoderm-like (EPL) cells - which lies between the naïve and primed states. EPL cells retain expression of pluripotency genes, upregulate primitive ectoderm markers, undergo a morphological change and have increased cell number.

These changes are facilitated by a complex signalling network hinging on the Mapk, Fgfr, Pi3k and mTor pathways. We use a factorial experimental design coupled with linear modelling and Bayesian regularised neural networks to understand which signalling pathways are involved in the transition between ESCs and EPL cells, and how they underpin changes in morphology, cell number, apoptosis, proliferation and gene expression. This approach allows for consideration of where pathways work antagonistically or synergistically.

Modelling showed that most properties were affected by more than one inhibitor, and each inhibitor blocked specific aspects of differentiation. These mechanisms underpin both progression of stem cells across the *in vitro* pluripotency continuum and serve as a model for pre-, peri- and post-implantation embryogenesis.

**Summary Statement:** L-proline acts as growth factor to modulate phosphorylation of the Mapk, Pi3k, Fgf and mTor signalling pathways to drive embryonic stem cells to primitive ectoderm-like cells.

## Introduction

Amino acids are present in the high micromolar to millimolar range in mammalian reproductive fluid (Aguilar and Reyley, 2005; Cetin *et al*., 2005; Harris *et al*., 2005) and are required to support normal embryo development *in vivo* (Van Winkle, 2001; Van Winkle *et al*., 2006; Bazer, Johnson and Wu, 2015). Consistent with this, supplementation of culture media with selected amino acids or certain groups of amino acids can be used to improve preimplantation development (Gardner and Lane, 1993; Lane and Gardner, 1997; Harris *et al*., 2005). For example, L-proline is a conditionally non-essential amino acid present in tubal fluid at ~140 μM in mice, ~150 μM in humans, ~100 μM in rabbits, 50-300 μM in sheep and ~200 μM in cows (Aguilar and Reyley, 2005; Cetin *et al*., 2005). *In vitro*, L-proline helps improve bovine oocyte maturation rates (Bahrami *et al*., 2019), promotes development to the blastocyst stage in the mouse system when added only during fertilization (Treleaven *et al*., 2021), and improves development when added to mouse embryo culture following fertilization (Morris *et al*., 2020).

In pluripotent mouse embryonic stem cells (ESCs), which are an *in vitro* model of mammalian embryo development, the addition of L-proline, either in purified form or as part of HepG2 conditioned medium (MEDII), stimulates differentiation to a second pluripotent population known as early primitive ectoderm-like cells (EPL cells) (Rathjen *et al*., 1999; Washington *et al*., 2010)); also known as proline-induced cells (PiCs) (Casalino *et al*., 2011; Comes *et al*., 2013; D’Aniello *et al*., 2015, 2017; Patriarca *et al*., 2021; Minchiotti *et al*., 2022)). EPL cells/PiCs are metastable, as they revert to naïve ESCs upon removal of L-proline (Rathjen *et al*., 1999; Washington *et al*., 2010; Casalino *et al*., 2011).

The transition to EPL cells recapitulates many of the features of the conversion of inner cell mass (ICM) cells in the 4.5 *days post coitum* (dpc) mouse embryo to pluripotent primitive ectoderm by ~5.5 dpc, with the primitive ectoderm now primed to gastrulate and form the 3 multipotent germ layers (Snow, 1977; Washington *et al*., 2010; Rivera-Pérez and Hadjantonakis, 2015). These similarities include the following: EPL cells are more primed to differentiate than ESCs, and represent a pluripotent population akin to formative pluripotency (Smith, 2017; Hoogland and Marks, 2021; Wang *et al*., 2021; Glover *et al*., 2022). The expression of the ICM marker *Rex1* in EPL cells is reduced, and there is increased expression of the primitive ectoderm markers *Fgf5* and *Dnmt3b* (Rathjen *et al*., 1999; Washington *et al*., 2010; Glover *et al*., 2022). Colonies undergo a change in morphology from round and domed to flattened monolayers with irregular borders, and cell-cycle time is reduced from 11 h to 8 h (Stead *et al*., 2002; Washington *et al*., 2010; Glover *et al*., 2022). The continued presence of L-proline in culture then drives EPL cells to neural cells by a series of embryologically relevant intermediate cell types (Rathjen *et al*., 2002; Shparberg *et al*., 2019).

In ESCs, L-proline is taken up via the Snat2 (*Slc38a2*) transporter (Tan *et al*., 2011). The mechanisms by which L-proline stimulates development/differentiation include (i) acute activation of signalling pathways, (ii) epigenetic remodelling and (iii) regulation of intracellular metabolism (Washington *et al*., 2010; Casalino *et al*., 2011; Comes *et al*., 2013; D’Aniello *et al*., 2015, 2017; Tan *et al*., 2016). Collectively, these mechanisms modify a range of emergent properties that drive developmental progression and differentiation (Washington *et al*., 2010) and are consistent with L-proline behaving as a growth factor (Morris *et al*., 2020).

The mTorc1 pathway is required for L-proline-mediated improvement in mouse preimplantation development and L-proline also activates the Erk1/2 and Akt pathways during this time (Morris *et al*., 2020). When added to ESCs, L-proline acutely activates the same signalling pathways (Lonic, 2006; Washington *et al*., 2010), as well as the p38 pathway (Tan *et al*., 2016) and selective inhibition of mTorc1 (with rapamycin) or Mek1/Erk1/2 (with U0126) or P38 (with SB203580 or PP2) prevents upregulation of the EPL cell marker *Dnmt3b* (Lonic, 2006; Washington *et al*., 2010). On the other hand, inhibition of the Pi3k/Akt pathway with LY294002 prevents the morphology change and increase in proliferation but allows the associated gene-expression changes to occur (Lonic, 2006). Thus, a number of signalling pathways are involved in the transition from ESCs to EPL cells, selective inhibition of pathways blocks different aspects of the transition, and collectively this shows that L-proline activates a complex signalling network. However, these experiments did not comprehensively measure changes in a range of emergent properties and marker expression to better understand this complex signalling.

To explore this, we employed inhibitors of Mek1, Fgf receptor (Fgfr), Pi3k, mTorc1 and P70-S6 kinase (S6k) alone and in all combinations. The results of these factorial experiments were analysed by linear, multiple interaction and Bayesian regularised neural network with a Gaussian prior (BRANNGP) modelling to generate complementary models that avoid issues of model overfitting (Woolf *et al*., 2005; Burden and Winkler, 2008; Winkler and Burden, 2012; Epa *et al*., 2013) and reveal synergistic and antagonistic effects. This approach is most commonly used to determine drug interactions (Sorokin *et al*., 2018; Julkunen *et al*., 2020; Panina *et al*., 2020) and is becoming increasingly used in stem cell biology (Chang and Zandstra, 2004; Prudhomme, Duggar and Lauffenburger, 2004; Audet, 2010; Jakobsen *et al*., 2014; Ireland *et al*., 2020).

## Results

### ESC-to-EPL cell transition alters gene expression and emergent properties

ESCs were maintained in either 330 or 1000 U/mL LIF and then directed to differentiate into EPL cells by addition of 400 μM L-proline for 6 days. In addition, cells were allowed to undergo spontaneous differentiation in the absence of LIF (**Fig. 1A**). ESCs grown in 330 or 1000 U/mL LIF maintained their dome-shaped colonies and showed no differences in morphology score. Colony morphology changed significantly in the presence of L-proline to flattened epithelial-like colonies, whereas cells allowed to spontaneously differentiate underwent a more robust morphology change (**Fig. 1A-B**), consistent with these cells undergoing differentiation beyond the EPL cell stage (Tan *et al*., 2011; Minchiotti *et al*., 2022).

**Figure 1.**
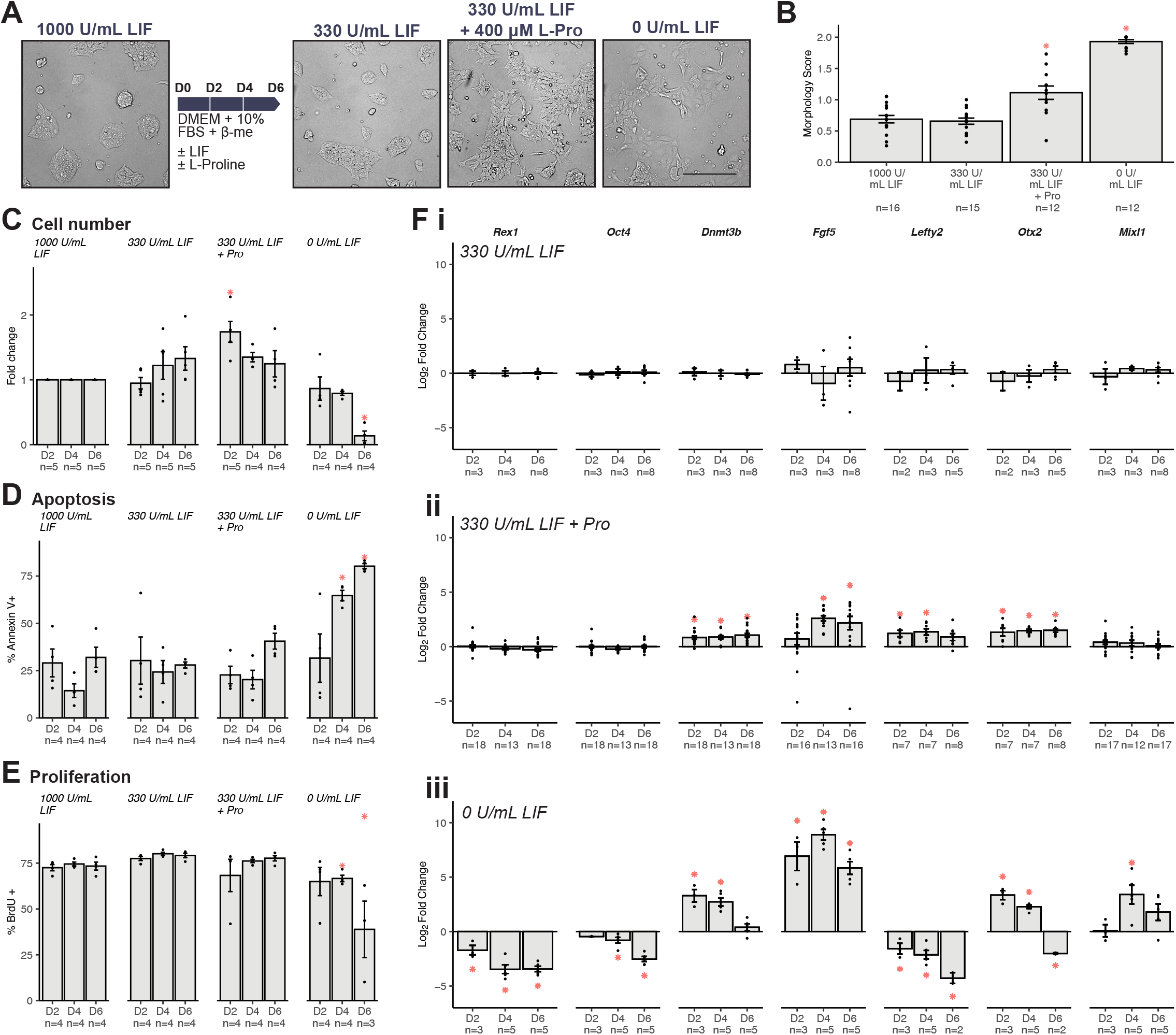
L-proline drives ESCs to EPL cells. A. Representative images showing ESCs that have self-renewed in medium containing 1000 U/mL LIF or differentiated in medium containing 330 U/mL LIF, 330 U/mL LIF + L-proline or no LIF for 6 days. Scale bar = 100 μm. **B.** Colony morphology was scored at day 6. Cell number (**C**), apoptosis (**D**) and proliferation (**E**) were measured at days 2, 4 and 6. **F.** At days 2, 4 and 6, changes in expression of pluripotency genes (*Rex1* and *Oct4*), primitive ectoderm markers (*Dnmt3b, Fgf5, Lefty2* and *Otx2*), and mesendoderm *genes (Mixl1*) in cells grown in medium containing (i) 330 U/mL LIF, (ii) 330 U/mL LIF + L-proline and (iii) no LIF or L-proline. All samples were normalised to *β-Actin* and then to cells grown in 1000 U/mL LIF. All graphs (B-F) show mean ± SEM with individual data points. Data were analysed using one-way ANOVA with Dunnett’s multiple comparisons test to cells grown in 1000 U/mL LIF, *\*P* < 0.05.

Cell number, apoptosis and proliferation were quantified at days 2, 4 and 6 of differentiation. Cell number was normalized to cells growing in 1000 U/mL LIF. Cells growing in 330 U/mL LIF + 400 μM L-proline increased cell number by 1.7-fold at day 2 (**Fig. 1C**), consistent with previous results for ESCs grown in 1000 U/mL LIF + L-proline (Washington et al., 2010). No significant changes in proliferation or apoptosis were observed (**Fig. 1D-E**). Cells undergoing spontaneous differentiation without LIF or L-proline were dying by day 6: Cell number was reduced by 7.8-fold (**Fig. 1C**), apoptosis increased 4.2-fold at day 4 and 2.3-fold at day 6 (**Fig. 1D**), and proliferation decreased 2.0-fold at day 6 (**Fig. 1E**), suggesting deficiencies in medium formulation and therefore a reduced capacity to support growth of differentiating cells.

After 6 days of differentiation, gene expression was profiled, focusing on pluripotency genes *Rex1* and *Oct4*, primitive ectoderm markers *Dnmt3b, Fgf5, Lefty2* and *Otx2*, and the mesendoderm marker *Mixl1*. There were no differences in the expression of any of the genes between ESCs grown in 330 or 1000 U/mL LIF (**Fig. 1Fi**). Cells grown in 330 U/mL LIF + 400 μM L-proline had comparable expression of *Rex* and *Oct4*, indicating maintenance of pluripotency, and the expression of the mesendoderm marker, *Mixl1*, also did not change. However, the expression of all primitive ectoderm markers increased (**Fig. 1Fii**). Cells allowed to spontaneously differentiate had a gene expression profile consistent with rapid, unregulated differentiation: Reduced expression of *Rex1* and *Oct4*, a wave of expression of *Dnmt3b, Fgf5*, and *Otx2* but with *Dnmt3b* expression returning to baseline and that of *Otx2* decreased by day 6. The expression of *Lefty2* remained strongly reduced throughout, while *Mixl1* expression increased significantly (**Fig. 1Fiii**).

### L-proline-mediated phosphorylation of signalling pathway intermediates

We examined the phosphorylation status of signalling pathway intermediates drawn from the Stat3, Fgf, Mek1/Erk1/2, Pi3k/Akt, mTor, p38 and Pkc pathways (**Fig. 2A**), each of which is known to play a role in pluripotence and/or differentiation of ESCs (Kunath *et al*., 2007; Lanner and Rossant, 2010; Washington *et al*., 2010; Cherepkova, Sineva and Pospelov, 2016; Tan *et al*., 2016).

**Figure 2.**
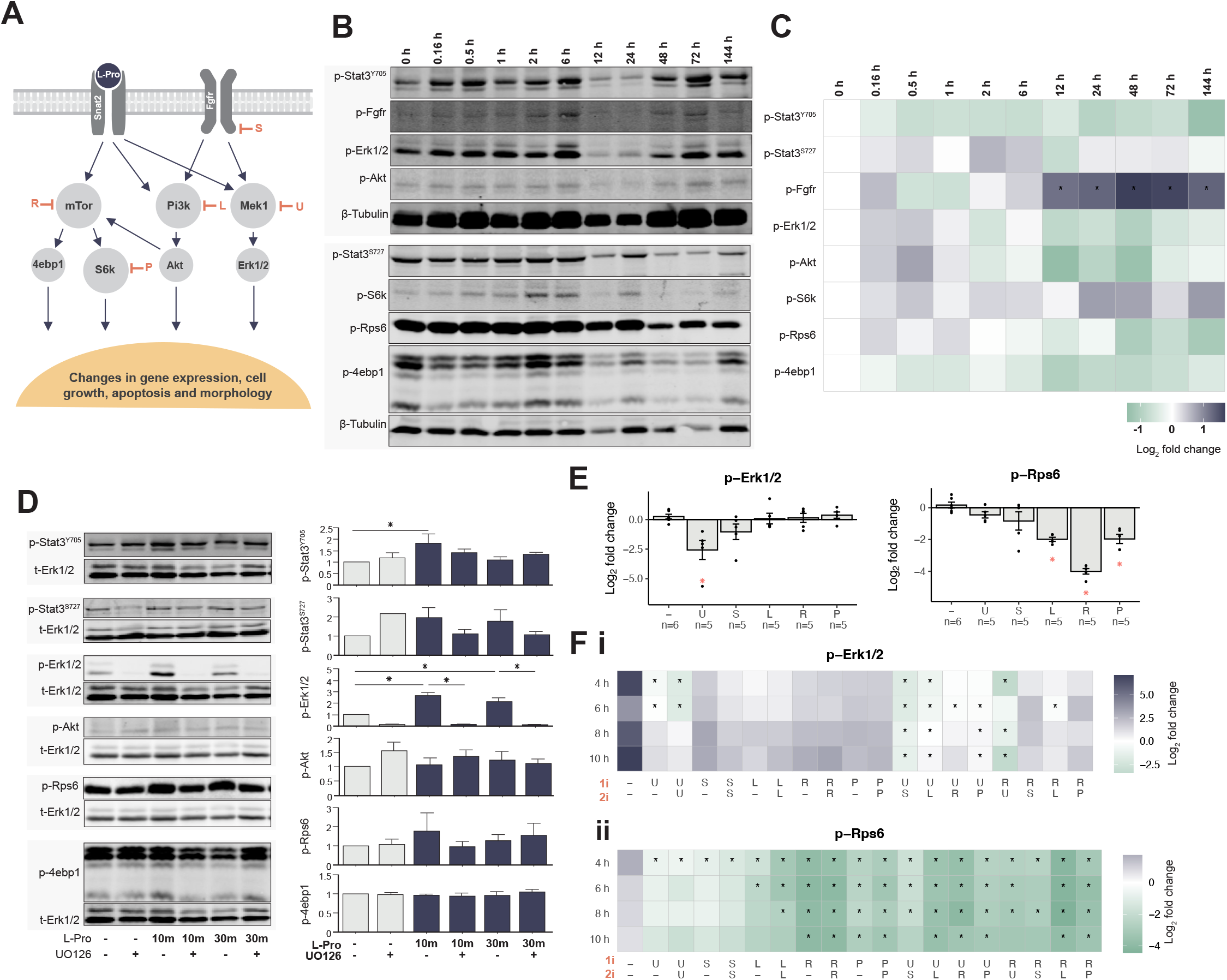
L-proline acts through Fgfr, Mapk, Pi3k and mTor signalling pathways. **A.** L-proline enters the cell via the SNAT2 transporter and activates the Mapk pathway (which can be inhibited by U0126, U), the Pi3k pathway (which can be inhibited by LY294002, L), the mTor pathway (which can be inhibited by rapamycin, R), the downstream mTor kinase S6k (which can be inhibited by PF-4708671, P), or indirectly activates the Fgfr (which can be inhibited by SU5402, S). Activation or inhibition of these pathways affects both gene expression and emergent cellular properties. **B.** Naïve ESCs were grown in medium containing 330 U/mL LIF + L-proline for up to 6 days (144 h). Cell lysates were analysed by western blotting for: p-Stat3^Y705^ or p-Stat3^S727^), p-Fgfr^Y653/Y654^, p-Erk1/2^T202/Y204^, p-Akt^S473^, p-S6k^T389^, p-Rps6^S235/S236^ and p-4ebp1^T37/T46^. Representative images shown. **C.** Quantification of western blot bands. **D.** Naïve ESCs were serum starved in DMEM + 0.1% FBS for 4 h, with 5 μM U0126 added for the final 30 min, where indicated. Cells were then left untreated or treated with 1 mM L-proline for 10 or 30 min. **E.** 400 μM L-proline and signalling pathway inhibitors were added to naïve ESCs in 1000 U/mL LIF. After 2 h, cell samples were analysed for p-Erk1/2 (**i**) and p-Rps6 (**ii**). **F.** ESCs were treated with 400 μM L-proline alone, or L-proline and a signalling pathway inhibitor (1i) at 0 h. At 2, 4, 6, or 8 h, a second dose of the same inhibitor or a different inhibitor (2i) was added and samples were collected 2 h later. Cell samples were analysed for p-Erk1/2 (**i**) and p-Rps6 (**ii**). For all western blot data band intensity was normalised to *γ*-Tubulin (C, E, F) or t-Erk1/2 (D) and normalized to an untreated ESC sample. Graphs show either fold change ± SEM (D) or log_2_ fold change ± SEM and individual data points (E). Heatmaps show mean log_2_ fold change. Data were analysed using one-way ANOVA with *post hoc* Tukey’s (D) or Dunnett’s (C, E, F) multiple comparison test, \**P* < 0.05.

Naïve, self-renewing ESCs were switched from 1000 U/mL LIF to EPL cell medium containing 330 U/mL LIF + 400 μM L-proline and the phosphorylation status of the pathway intermediates was quantified by western blot over the short (0-12 h) and long term (1-6 days). Of these, only phosphorylation of Fgfr increased significantly (from 12 h onwards) (**Fig. 2B-C**). The P38 and Pkc pathways, including the downstream Hsp27, which had previously been shown to be altered by addition of L-proline (Tan *et al*., 2016), were not detected in ESCs ± L-proline, under our conditions (**Fig. S1**). L-proline acutely increased phosphorylation of Erk1/2 and Stat3^Y705^ in these conditions (**Fig. 2D**).

### Signalling pathway inhibition illustrates pathway cross-talk

The effect of signalling pathway inhibitors, alone or in combination, on L-proline-mediated pathway activity was assessed using the following: (i) Mapk pathway using the Mek1/2 inhibitor U0126 (U) (Favata *et al*., 1998); (ii) Fgfr pathway using receptor antagonist SU5402 (S) (Mohammadi *et al*., 1997); (iii) Pi3k/Akt pathway using the Pi3k inhibitor LY294002 (L) (Vlahos *et al*., 1994); (iv) mTorc1 pathway using the mTorc1 complex inhibitor rapamycin (R) (Sabers *et al*., 1995); (v) mTorc1 pathway at the downstream kinase, S6k, using the inhibitor PF-4708671 (P) (Pearce *et al*., 2010).

An initial dose of U0126 suppressed L-proline-mediated Erk1/2 (downstream of Mek1) phosphorylation only for 6 h, after which phosphorylation returned to that seen with L-proline only (**Fig. 2E, Fi**). Furthermore, any second dose of U0126 failed to suppress Erk1/2 phosphorylation for longer than 6 h (**Fig. 2Fi**), suggesting Erk1/2 phosphorylation was no longer controlled by Mek1.

None of the other 4 pathway inhibitors by themselves suppressed Erk1/2 phosphorylation on their own (**Fig. 2Fi**). However, extended suppression of Erk1/2 phosphorylation occurred up to 10 h when SU5401 or LY294002 or PF-4708671 were added after the initial addition of U0126 (**Fig. 2Fi**), suggesting Erk1/2 phosphorylation was now controlled by crosstalk involving an Fgfr/Pi3k/S6k axis.

Rps6 phosphorylation shows similar interplay between pathways. pRps6 phosphorylation was suppressed for up to 10 h by rapamycin and LY294002 (0.26 ± 0.02 SEM, **Fig. 2E**), consistent with it lying on the mTorc1/S6k signalling axis (**Fig. 2A**). However, its phosphorylation was also temporarily suppressed by inhibition of Mek1, Fgfr, and Pi3k, or combinations of Mek1 inhibition followed by Fgfr inhibition, or Mek1 inhibition followed by Pi3k inhibition (**Fig. 2Fii**). These results point to complex, dynamic changes in signalling pathway activity over time in the presence of L-proline.

### Factorial experiments reveal relationships between emergent properties

To further our understanding of pathway interactions and their effect of emergent properties, a factorial experiment was designed for the differentiation of ESCs to EPL cells over 6 days in the presence of all possible combinations of the five inhibitors (**Fig. 3A**). On days 2 and 4, cells were counted, replated, and samples were collected to quantify apoptosis and cell proliferation by flow cytometry. At day 6, in addition to measurements of apoptosis and cell proliferation, cells were imaged for colony morphology and cell number, and qPCR was used to quantify changes in the expression of marker genes.

**Figure 3.**
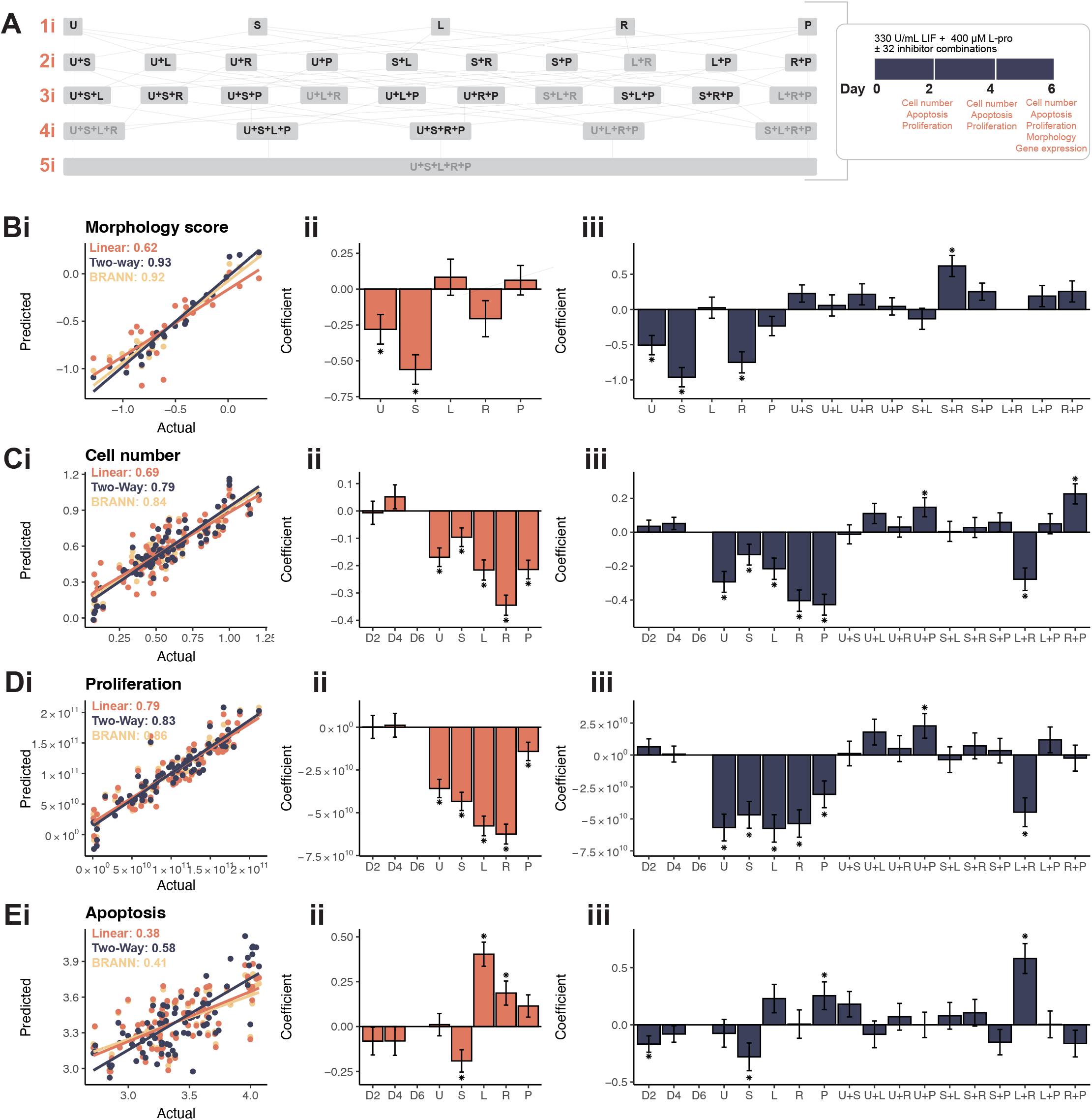
Signalling pathway inhibitors regulate emergent properties during the ESC-to-EPL cell transition. **A.** Cartesian product experimental design showing all combinations of the five signalling pathway inhibitors. Naïve ESCs were differentiated over 6 days in 330 U/mL LIF + 400 μM L-proline with combinations of five inhibitors (U: U0126; S: SU5402; L: LY294002; R: rapamycin; P: PF-4708671). Colony morphology was scored on day 6 (**B**), while cell number (**C**), proliferation (**D**) and apoptosis (**E**) were recorded at days 2, 4 and 6. Data was averaged across biological replicates where n ≥ 3. To correct for non-normal distributions, morphology and apoptosis data were log transformed, and proliferation data was raised (x^6^). (**i)** Data was modelled using either linear modelling, linear modelling with two-way interaction terms or a Bayesian regularised neural network (BRANNGP). Fit of each model is shown comparing the actual fit with the prediction from the model. (**ii)** Coefficients for each variable ± SEM for standard linear model. (**iii)** Coefficients for each variable ± SEM for linear model with two-way interaction terms. Significance is denoted as \**P* < 0.05.

At day 2, the 8 conditions which contained both LY294002 and rapamycin had poor viability: Cell number had reduced by 90%, apoptosis had increased by more than 50%, and proliferation reduced by nearly 40% (**Fig. S2**). By day 4, very few cells were present. These conditions were considered to be non-viable, and were not considered in the day 4 and 6 measurements.

To get a broad understanding of relationships between emergent properties and gene expression, a correlation matrix was generated using all data from all viable inhibitor combinations (**Fig. S3A**). Expected correlations were observed, such as (i) the positive correlation between cell number and proliferation, (ii) the negative correlation between cell number and apoptosis across the 6 days of transition to EPL cells, and (iii) the coupling of expression of pluripotency markers *Oct4* and *Rex1* and EPL-cell markers *Dnmt3b* and *Fgf5*. However, more nuanced correlations were observed, including positive correlation (i) between proliferation (from day 4 onwards) and morphology, and (ii) between proliferation and expression of differentiation-related genes (*Dnmt3b*, *Fgf5*, and *Mixl1* at days 4 and 6). A high correlation was also observed between expression of the primitive ectoderm marker *Lefty2* and the pluripotency markers *Oct4* and *Rex1*.

### Results of data modelling help deconvolute complex signalling networks

To understand which signalling pathways or pathway combinations drive changes in gene expression and emergent properties, we generated multiple linear regression (MLR) and BRANNGP models (**Fig. 3B-Ei, Fig. 4i**). The coefficients underlying the fit of each model were used to indicate the direction and extent each inhibitor contributes to the response (**Fig. 3B-Eii, Fig. 4ii**). We also ran MLR with two-interaction terms to determine if inhibitors were acting independently (*additive), synergistically* or *antagonistically*. For models with interaction terms, the coefficients for the inhibitors alone are added to the coefficient for the interaction term. An *additive* effect is seen where there are significant effects for the inhibitors alone but no significant interaction effect. A *synergistic* effect is seen where there are significant effects for two inhibitors alone, and a significant interaction effect in the same direction. An *antagonistic* effect is seen where there are significant effects for two inhibitors alone, and a significant interaction effect in the opposite direction.

**Figure 4.**
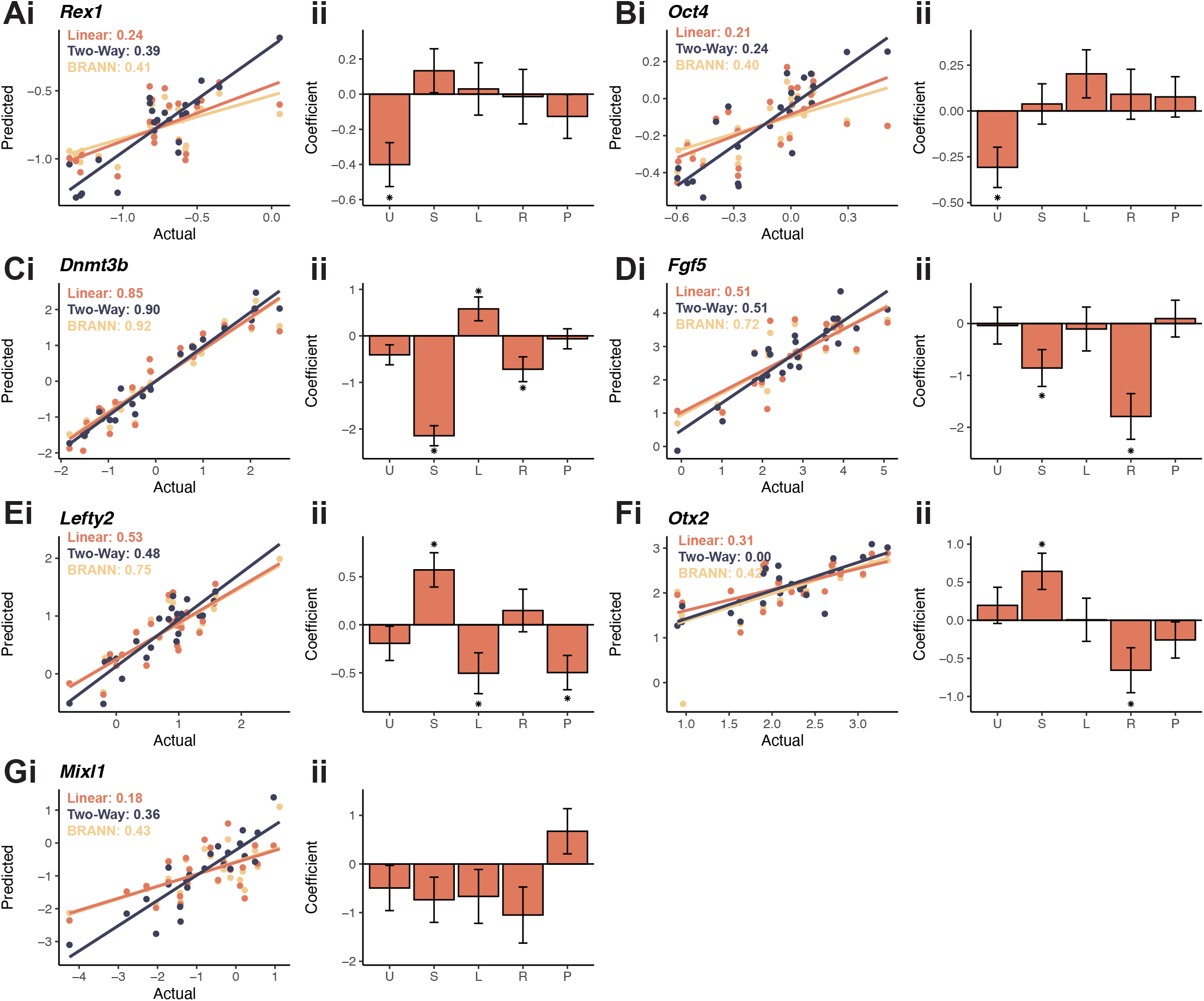
Signalling pathway inhibitors regulate gene expression during the ESC-to-EPL cell transition. Naïve ESCs were differentiated over 6 days in 330 U/mL LIF + 400 μM L-proline with combinations of five inhibitors (U: U0126; S: SU5402; L: LY294002; R: rapamycin; P: PF-4708671). At day 6, changes in expression of pluripotency genes (*Rex1*, **A** and *Oct4*, **B**), primitive ectoderm markers (*Dnmt3b*, **C**, *Fgf5*, **D**, *Lefty2*, **E** and *Otx2*, **F**), and mesendoderm *genes (Mixl1*, **G**). All samples were normalised to *β*-Actin and then to cells grown in 1000 U/mL LIF. Data was averaged across biological replicates where n ≥ 3. **(i)** Data was modelled using either linear modelling, linear modelling with two-way interaction terms or a Bayesian regularised neural network (BRANNGP). Fit of each model is shown comparing the actual fit with the prediction from the model. (**ii**) Coefficients for each variable ± SEM for standard linear model.

MLR fits with a higher adjusted R^2^ and a lower standard error (σ) value generally denote a better fit to the data (**Table S1**). An F-test was used to determine if MLR with interaction terms resulted in a significantly improved fit, taking into account the increased number of parameters (**Table S2**).

BRANNGPs were utilised to provide robust, potentially nonlinear models that may better explain structure-activity relationships without issues around overfitting and overtraining (Burden & Winkler, 2008; Winkler & Burden, 2012). Most data sets had improved R^2^ values when modelled using BRANNGP (**Table S1**). The average R^2^ value for BRANNGPs was 0.64 ± 0.07 compared to 0.48 ± 0.07 for MLR models or 0.53 ± 0.09 for MLR models with two-way interaction terms. Only apoptosis and proliferation models had R^2^ values for BRANNGP models similar to the MLR models. BRANNGP models with improved fit suggest additional nonlinear factors are involved. However, the nature and magnitude of these factors cannot as yet be deconvoluted from the BRANNGP models.

### Morphology is regulated by Erk1/2, Fgfr and mTor

The morphology data set (**Fig. S2**) was used to train standard MLR, MLR with two-way interaction terms, and BRANNGP models (**Fig. 3Bi**). The standard MLR model shows that addition of SU5402 or U0126 prevented changes in colony morphology normally expected in the presence of L-proline (**Fig. 3Bii**), resulting in cells which retained a domed, ESC-like appearance. An F-test indicated that MLR with two-way interaction terms provided a better fit than the MLR (**Table S2**). This improved model showed that SU5402, U0126 and rapamycin all prevent morphology change (**Fig. 3Biii**). There was no significant interaction between SU5402 and U0126 indicating that this effect was largely additive. There was a significant interaction between SU5402 and rapamycin mediating morphology change. This interaction coefficient was reversed, indicating an antagonistic effect.

### All inhibitors decrease cell number and proliferation

Modelling was performed on the cell number and proliferation data from all inhibitor combinations. MLR produced robust fits for both cell number and proliferation with R^2^ of 0.69 and 0.79 respectively (**Fig. 3C-Di**). All inhibitors significantly reduced cell number and proliferation, with rapamycin having the largest effect (**Fig. 3C-Dii**).

For both cell number and proliferation, the MLR with two-way interactions had an improved R^2^ (0.79 and 0.83 respectively), and the F-test showed significant improvement (**Table S2**). The individual effects were retained for each inhibitor, but multiple interaction effects were noted: i) U0126 and PF-4708671 were antagonistic in both cell number and proliferation models; ii) antagonism between rapamycin and PF-4708671 in the cell number model; iii) LY294002 and rapamycin were strongly synergistic for both cell number and proliferation models (**Fig. 3C-Diii**). An alternate model, which attempts to overcome non-normality of the proliferation input data by dividing the data into octiles, exhibited a very similar results to the linear model (**Fig. S4**).

### Apoptosis is differentially altered by each inhibitor

Apoptosis data generated an adequate fit using the MLR (R^2^ of 0.38), but this was significantly increased using the MLR with two-way interactions (R^2^ of 0.58 and F-test *P* < 0.05, **Fig. 3Ei, Table S2**). In the standard MLR model, SU5402 reduced apoptosis and LY294002 and rapamycin increased apoptosis (**Fig. 3Eii**). In the MLR with two-way interactions the individual effects between LY294002 and rapamycin were lost, and instead there was a strong synergistic effect between these inhibitors. The day 2 parameter was also significant. In conjunction with the reduction in proliferation, this explains the early cellular lethality of the combination of LY294002 and rapamycin, which does not affect the inhibitors when used alone. The apoptosis model retained the significant reduction in apoptosis with SU5402, and also showed that apoptosis was increased with PF-4708671 (**Fig. 3Eiii**). While both SU5402 and PF-4708671 reduce apoptosis, the reduction in proliferation (Fig. 3Cii) resulted in a net decrease in cell number (Fig. 3Dii). U0126 did not affect apoptosis in the presence of L-proline (**Fig. 3Eii**), indicating that the decrease in cell number elicited by U026 is due entirely to a decrease in proliferation (**Fig. 3Cii, Dii**).

### Gene expression is regulated by intracellular signalling

Modelling was also performed to assess how inhibitors impacted gene expression at day 6 (**Fig. 4, S5**). MLR for *Dnmt3b, Fgf5* and *Lefty2* had a robust fit (R^2^ > 0.5, **Fig. 4C-Ei**). Models for *Rex1, Oct4, Mixl1* and *Otx2* expression had poor to modest fit (R^2^ of 0.24, 0.21, 0,18 and 0.31 respectively, **Fig. 4A-Bi, F-Gi**). None of these models were significantly improved by switching to two-way interaction effect models (F > 0.05, **Table S2, Fig. S6).**

The MLR fit showed that U0126 decreased expression of the pluripotency genes *Rex1* and *Oct4* (**Fig. 4Aii, Bii**). The decrease was modest when compared to the strong downregulation of these genes for cells undergoing spontaneous differentiation (**Fig. 1Fiii**). There was significant U0126 and SU5402 interaction in the *Rex1* MLR model with interaction terms, suggesting some antagonism (**Fig. 4Aiii**).

Inconsistent changes in expression patterns occurred frequently for the four EPL markers – *Dnmt3b*, *Fgf5*, *Lefty2* and *Otx2*. For example, SU5402 decreased expression of *Dnmt3b* and *Fgf5* (**Fig. 4C-Dii**) but increased expression of *Lefty2* and *Otx2* (**Fig. 4E-Fii**). LY294002 increased *Dnmt3b* expression (**Fig. 4Cii**) but reduced that of *Lefty2* (**Fig. 4Eii**). Rapamycin decreased *Dnmt3b, Fgf5* and *Otx2* expression (**Fig. 4C-Dii, Fii**) but did not alter that of *Lefty2*. PF4708671 only reduced *Lefty2* expression (**Fig. 4Eii**). These results highlight that the expression of individual genes associated with the identity of EPL cells are associated with complex signalling pathway control. In all cases, 2 or 3 of the tested pathways regulated expression of each gene. The MLR models for the mesendoderm gene *Mixl1* were not robust enough to make strong biological statements (R^2^ of 0.18, **Fig. 4Gi**).

### Functional assay establishes how inhibitor treated cells fall on the pluripotency continuum

This lack of consensus in the inhibitors driving expression of EPL marker genes was further exemplified by data showing that many inhibitor combinations suppressed some but not all primitive ectoderm genes (**Fig. S5**). To address this, we ran a functional assay to identify the pluripotency capacity of each inhibitor combination: After 6 days of differentiation in the presence of L-proline and the various inhibitors, cells were allowed to spontaneously differentiate as embryoid bodies (EBs). Samples were collected on days 2, 3 and 4 and qRT-PCR was used to quantify expression of the primitive streak marker *Brachyury*. In the absence of any inhibitors, cells which were more naïve, like ESCs, took 4 days to upregulate *Brachyury*, compared to more primed cells, like EPL cells, which upregulated expression of *Brachyury* at day 2 (**Fig. S7A**). Across all inhibitor treated conditions, conditions which contained U0126 or LY294002 tended to upregulate *Brachyury* expression earlier, and conditions which contained rapamycin tended to upregulate *Brachyury* later (**Fig. 7Bii**). We also assessed the correlations between the slope of *Brachyury* upregulation to the other genes. Significant positive correlations were noted between *Brachyury* upregulation and *Dnmt3b, Fgf5* and *Mixl1*, but not the pluripotency markers *Rex1* and *Oct4*, or the more recently adopted primitive ectoderm markers *Lefty2* and *Otx2* (**Fig. S3B**).

## Discussion

### Signalling pathways active during ESC differentiation to EPL cells

This study used small molecule inhibitors to help elucidate the role of various signalling pathways that mediate self-renewal, differentiation, and other emergent properties such as colony morphology, cell number, proliferation, and apoptosis during the transition of ESCs to EPL cells (**Fig. 5A**); *viz* Mapk (using the Mek1 inhibitor U0126), Fgfr (using the antagonist SU5402), Pi3k (using the Pi3k inhibitor LY294002) and mTor pathways (using the mTorc1 complex inhibitor rapamycin, or the S6k inhibitor PF-4708671). These signalling pathways are acutely activated in response to L-proline (**Fig. 2D**) or have been previously associated with L-proline-mediated differentiation to EPL cells (Lonic, 2006; Washington *et al*., 2010; Tan *et al*., 2016).

**Figure 5.**
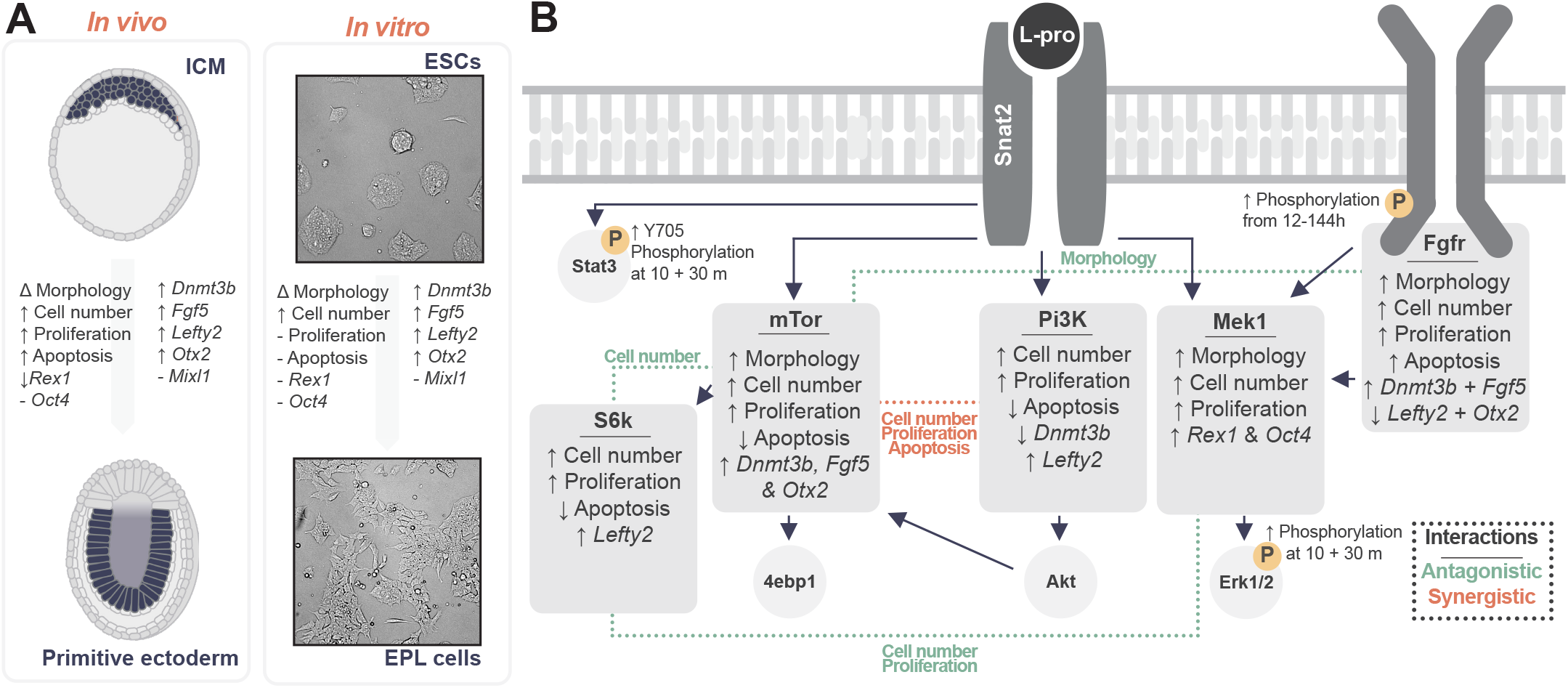
Summary of signalling pathway-mediated changes in emergent properties and gene expression during the ESC-to-EPL cell transition. **A.** The L-proline mediated ESC-to-EPL cell transition recapitulates the transition from the inner cell mass (ICM) to the primitive ectoderm (Snow, 1977; Coucouvanis and Martin, 1999; Brennan *et al*., 2001; Hart *et al*., 2002; Pelton *et al*., 2002; Watanabe *et al*., 2002; Acampora, Di Giovannantonio and Simeone, 2013). **B.** Results of linear modelling show that L-proline acts through each signalling pathway to control different aspects of differentiation. Interaction effects were noted between some inhibitor combinations, and these are shown in dotted lines. Red dotted lines show synergistic effects where two inhibitors produce a response larger than either alone. Green dotted lines show antagonistic effects where two inhibitors produce a response less than the sum of either alone.

In the absence of inhibitors, L-proline increased pathway phosphorylation (**Fig. 5B**), including acute phosphorylation of Stat3^Y705^ and Erk1/2 within 10 min (**Fig. 2D-E**), which suggests L-proline rapidly induces changes in pathways known to be important for maintenance/loss of pluripotency (Stavridis, Collins and Storey, 2010; Huang *et al*., 2014). Over the course of differentiation, Fgfr phosphorylation increased but with no change in the canonical intermediate Erk1/2 (**Fig. 2B-C**), suggesting Fgfr is likely signalling through other intermediates such as Pkc, Pi3k, Src, Stat1, P38 and Jnk (Dailey *et al*., 2005).

When signalling pathway inhibitors were used in the presence of L-proline, signalling pathway cross talk led to maintenance of Mapk signalling: Erk1/2, immediately downstream of U0126 target Mek1, had decreased phosphorylation in the presence of this inhibitor but only out to 6 h. A second dose of U0126 did not extend this time (**Fig. 2Fi**). This effect is not unique to U0126: Erk1/2 phosphorylation is only transiently reduced when a variety of Mek1 inhibitors (PD98059, PD184352, PD0325901 and U0126) is added to the culture medium of ESCs (Chen *et al*., 2015). Together these results indicate it’s unlikely the reduced Erk1/2 phosphorylation in the presence of U0126 is due to loss of inhibitor activity but rather due to Erk1/2 phosphorylation now being maintained by pathway cross-talk, which bypasses Mek1. Since reduced phosphorylation of Erk1/2 in the presence of U0126 could be extended out to 10 h by also including the Fgfr inhibitor SU5402 or the Pi3k inhibitor LY294002, one possibility is that the Fgfr-Pi3k-Akt axis now sustains L-proline-mediated phosphorylation of Erk1/2 (Dailey *et al*., 2005). This complex network with multiple inputs speaks to the importance of Erk1/2 signalling to avoid widespread apoptosis, as seen in *Erk1^-/-^/Erk2^-/-^* ESCs (Chen *et al*., 2015).

### Modelling reveals inhibitors which alter aspects of the transition of ESCs to EPL cells

We designed a factorial study to assess how signalling pathways influence a variety of properties during the L-proline-mediated transition from ESCs to EPL cells (**Fig. 5**). No single inhibitor was sufficient to explain all the changes in gene expression and emergent properties during ESC differentiation to EPL cells. Rather, our modelling suggests that these signalling pathways have discrete roles within this transition, likely supported by signalling pathway crosstalk.

From the combinational experiments, we note the following for each pathway inhibitor:

i. When the Mapk/Erk1/2 pathway was inhibited by U0126, cells didn’t undergo the morphological change associated with the presence of L-proline (**Fig. 3Aii**) even though expression of *Rex1* and *Oct4* was decreased (**Fig. 4A-B, Fig. 5**). The decrease in *Rex1* and *Oct4* was less than cells undergoing spontaneous differentiation (**Fig. 1Fiii**) but indicates disruption of the pluripotency gene regulatory network (Kim *et al*., 2008).
ii. When the Fgfr was inhibited by SU5402, cells again didn’t undergo a morphology change (**Fig. 3Bii**) but instead the expression of the EPL-cell markers, *Dnmt3b* and *Fgf5*, which is increased in the presence of L-proline alone, was blocked. In contrast, expression of EPL-cell markers *Otx2* and *Lefty2* were increased in the presence of this inhibitor (**Fig. 4C-F, Fig. 5**). This suggests that Fgfr inhibition at least partially blocks the transition.
iii. When the Pi3K/Akt pathway was inhibited with LY294002, the L-proline-mediated change in colony morphology was still permitted, as was the increased expression of EPL-cell markers, *Dnmt3b* and *Fgf5*. The L-proline-mediated increase in *Otx2* expression was also allowed but the L-proline-mediated increase in *Lefty2* expression was suppressed. An early increase in *Lefty2* expression is associated with the transition of ESCs to EPL cells and reduction in expression as pluripotency is lost (Harvey *et al*., 2010) but these results suggest increased *Lefty2* expression is not obligatory for the transition.
iv. When mTorc1 was inhibited by rapamycin, ESCs underwent the L-proline-mediated change in morphology but the increased expression of *Dnmt3b, Fgf5* and *Otx2* was suppressed. This suppressed gene expression confirms previously published data (Washington *et al*., 2010). but in that work rapamycin also blocked the morphology change, which we did not observe. Earlier protocols generated EPL cells using 1000 U/mL LIF and L-proline, which inconsistently upregulated expression of the primitive ectoderm marker *Fgf5* (Washington *et al*., 2010). Here, we reduced LIF to 330 U/mL LIF, which results in robust upregulation of *Fgf5* expression (Harvey *et al*., 2010; Glover *et al*., 2022). These results highlight the sensitive balance between the cytokine LIF and the growth-factor-like properties of L-proline in promoting directed differentiation.
v. When the S6k branch of the mTorc1 pathway was inhibited with PF-4708671 it, like rapamycin, failed to prevent the L-proline-mediated change in colony morphology but unlike rapamycin it did not suppress the L-proline-mediated increase in the expression of the EPL-cell markers *Dnmt3b*, *Fgf5* and *Otx2* (**Fig. 4C-D, F, Fig. 5)**. This suggests that L-proline’s stimulation of expression of these markers requires the 4ebp1 branch of the mTorc1 pathway.

All 5 inhibitors reduced cell number and reduced the rate of proliferation compared to L-proline (**Fig. 3Cii, Dii**) but different effects were seen on apoptosis (Fig. 3Eii). Neither inhibition of the Mek/Erk1/2 pathway with U0126 nor S6k with PF-4708671 affected apoptosis, indicating that reduced cell numbers result from reduced proliferation. However, inhibition of mTorc1 with rapamycin increased apoptosis, supporting a role that mTorc1 signalling via the 4ebp1 branch is anti-apoptotic (Nawroth *et al*., 2011; Pons *et al*., 2011; Yellen *et al*., 2011). This branch is also pro-proliferative, and may explain why rapamycin compromises proliferation more than PF-4708671 (Dowling *et al*., 2010; Nawroth *et al*., 2011) (**Fig. 3Dii**). LY294002 led to an increase in apoptosis in addition to the decrease in proliferation, which is in line with Pi3k as a strong mediator of cell survival and progression (Chang *et al*., 2003; Takahashi, Murakami and Yamanaka, 2005; Tsurutani *et al*., 2005; Yu and Cui, 2016). Fgfr signalling produces cell- and state-specific effects on apoptosis and proliferation (Dailey *et al*., 2005), and reduced apoptosis observed with SU5402 provides further evidence for this. Collectively, these results highlight biological system complexity and make it difficult, if not impossible, *a priori* to determine outcomes even when a single inhibitor is used.

### Primitive ectoderm markers reflect spatial and temporal contributions to the EPL-cell transition

We selected four primitive ectoderm markers to assess how cells transitioned to EPL cells: *Dnmt3b, Fgf5, Lefty2* and *Otx2*. We found that these primitive ectoderm genes had similar expression patterns in differentiation to EPL cells (**Fig. 1F**) but behaved contrarily in inhibitor treated conditions (**Fig. 5B**).

In standard culture conditions, L-proline treated cells had significantly increased expression of *Dnmt3b* and *Otx2* across days 2 to 6 and *Fgf5* at days 4 and 6, whereas *Lefty2* expression was transiently increased at days 2 and 4 (**Fig. 1Fii**). Two of the inhibitors – SU5402 and LY294002 – has gene expression profiles which were less straightforward: cells treated with SU5402 had decreased *Dnmt3b* and *Fgf5* and increased *Lefty2* and *Otx2*, and cells treated with LY294002 had increased Dnmt3b expression but decreased *Lefty2* expression (**Fig. 4, 5**).

This difference may be due to different gene functions (i.e. *Dnmt3b* as a methyltransferase), or may reflect temporal or spatial expression patterns. We measured gene expression at day 6, which likely missed the transient peak of *Lefty2* expression as seen in our data (**Fig. 1Fii**), and in previous studies using EPL cells derived from embryoid bodies MEDII which also showed transient upregulation of *Lefty2* from days 1 to 4 (Harvey *et al*., 2010). This also explains why there were negative correlations between *Dnmt3b* and morphology changes (**Fig. S3A**). The return to baseline of *Lefty2* also makes sense considering significant positive correlations with stably-expressed pluripotency genes *Rex1* and *Oct4* (**Fig. 1Fii, S3A**).

To help assess ambiguity between markers, we included a functional assay which measured *Brachyury* expression as cells underwent spontaneous differentiation (**Fig. 7A**). We noted that conditions containing LY294002 tended to upregulate Brachyury earlier than average (**Fig. 7B**), suggesting that they were further along the pluripotency continuum and more like EPL cells. Conditions containing SU5402 tended to upregulate expression of *Brachyury* after 3 days, placing them in the middle of the continuum. Rapamycin tended to produce the naivest cells, though this may be skewed by lack of data from the non-viable conditions. No significant changes in expression of *Mixl1* were observed either in the absence or presence of the inhibitors (**Fig. 1Fii, 4G**), indicating that cells did not form mesendoderm.

### Modelling reveals synergy and antagonism in emergent properties

Biological complexity is further highlighted when two or more inhibitors were used together. Two-way interaction effects were used to determine if these pathways were independent (no interaction effects), antagonistic (where blocking two pathways simultaneously leads to a dampened effect compared to the sum of the two inhibitors individually) or synergistic (where blocking two pathways simultaneously leads to an increased effect compared to the sum of the two inhibitors individually). The emergent property results, but not the gene expression results, produced models which indicated interactions between pathways (**Table S2**).

Antagonistic effects were noted for Mek1 and S6k, where the combination of inhibitors U0126 and PF-4708671 attenuated the inhibition of both cell number and proliferation compared with the use of each of the inhibitors alone (**Fig. 3C-Diii, Fig. 5**); the combination of mTor and S6k inhibitors (rapamycin and PF-4708671) attenuated the inhibition of cell number (**Fig. 3Ciii**); and the combination of inhibitors for Fgfr and mTor (SU5402 and rapamycin) promoted the change in colony morphology, which the individual inhibitors prevented (**Fig. 3Biii**). These pathways likely coalesce on common downstream intermediates or transcription factors, or suppress other pathways through cross-talk (Mendoza, Er and Blenis, 2011; Aksamitiene, Kiyatkin and Kholodenko, 2012; Wang *et al*., 2013; Arkun, 2016).

Addition of both LY294002 and rapamycin resulted in strong synergistic effects that reduce cell number and proliferation and increase apoptosis (**Fig. 3C-Eiii, Fig. 5**) resulting in non-viable cells. Both pathways have been shown to individually reduce proliferation and increase apoptosis (Fingar *et al*., 2002; Jirmanova *et al*., 2002; Murakami *et al*., 2004; Gross, Hess and Cooper, 2005), and result in large defects in cell survival when used in combination in T cells, glioma cells, and small cell lung cancer cells (Breslin *et al*., 2005; Takeuchi *et al*., 2005; Tsurutani *et al*., 2005).

### Understanding L-proline mediated signalling in early embryogenesis

We have shown that L-proline activates several signalling pathways including the Mapk, Fgfr, Akt and mTor pathways to facilitate the transition of ESCs to EPL cells (**Fig. 5**). While changes in cell signalling are generally though to initiate changes in cell function, it is possible that other mechanisms of L-proline-mediated differentiation, including metabolic flux and epigenetic changes, may alter the cellular landscape to facilitate further changes in cell signalling. This has been seen previously with autocrine Fgf4 activation of the Fgfr as cells undergo differentiation (Kunath *et al*., 2007).

The L-proline-mediated transition of ESCs to EPL cells demonstrates the progression of cells from a naïve to primed state in the pluripotency continuum (D’Aniello *et al*., 2017; Morgani, Nichols and Hadjantonakis, 2017), and recapitulates aspects of peri- and post-implantation embryogenesis. The results are consistent with other growth factor-like role for L-proline including facilitating preimplantation embryo development (Morris *et al*., 2020; Treleaven *et al*., 2021), and differentiation of pluripotent cells towards neuroectoderm (Rathjen *et al*., 1999, 2002; Pelton *et al*., 2002; Harvey *et al*., 2010; Washington *et al*., 2010; Shparberg *et al*., 2019). L-proline mediated differentiation provides a useful model for studying embryonic development *in vitro*.

## Methods

### Cell culture

All cell culture was performed at 37°C, 5% CO_2_ in a humidified incubator. D3 ESCs (Doetschman *et al*., 1985) were maintained in ESC self-renewal medium containing DMEM (Sigma), 10% FBS (AusGeneX), 1000 U/mL LIF (Neuromics), 0.1 mM β-merceptoethanol (β-Me; Sigma) and Pen/Strep consisting of 50 U/mL penicillin (Sigma), and 50 μg/mL streptomycin (Sigma). Cells were grown as a monolayer, and passaged using Trypsin-EDTA (Sigma), and replated at 2,000-20,000 live cells/cm^2^ (Glover *et al*., 2022).

ESCs were differentiated to EPL cells by culturing 20,000 cells/cm^2^ in EPL cell medium (90% DMEM, 10% FBS, Pen/Strep, 0.1 mM β-Me, 330 U/mL LIF, 400 μM L-proline, Sigma) for 6 days, with passage every two days. As controls, ESCs were also cultured for 6 days with LIF reduced to 330 U/mL LIF, or allowed to spontaneously differentiate in without LIF or L-proline (Glover *et al*., 2022).

The effect of signalling pathway inhibitors (alone and in combination) on the transition of ESCs to EPL cells was tested (**Fig. 2E-F**). The inhibitors were as follows: Mek1 inhibitor U0126 (U, 5 μM, Selleck); Fgfr inhibitor SU5402 (S, 5 μM, MedChem Express); Pi3k inhibitor LY294002 (L, 5 μM, Selleck); mTorc1 inhibitor rapamycin (R, 10 nM, Selleck) and S6k inhibitor PF-4708671 (P, 10 μM, MedChem Express). All inhibitors were solubilized in DMSO, and a vehicle control containing the maximum concentration (0.22%) of DMSO was included.

At days 2, 4 and 6, differentiating ESCs treated with 1000 u/mL LIF or 330 U/mL LIF, and ESCs treated with L-proline ± inhibitor(s) were analysed for 3 emergent properties (cell number, apoptosis, and proliferation), as well as phosphorylation of various signalling pathway intermediates. Cells at day 6 were also analysed for colony morphology, changes in gene expression, and differentiation potential, as described below. Data were collected over 5 independent experiments.

### Measurement of cell number and colony morphology

Cell counts were measured with a haemocytometer following the addition of 0.4% Trypan Blue solution (Glover *et al*., 2022) to a single-cell suspension obtained following trypsinisation.

Colony morphology was quantified based on images collected from an Olympus IX-81 inverted microscope. Images were deidentified and colony morphology scored based on a predetermined scale: Round, domed (ESC) colonies were scored as 0. Flat, irregular, partially differentiated colonies were scored as 1, and fully differentiated colonies consistent with EPL cells were scored as 2 (Glover *et al*., 2022). Scoring was performed on all colonies (10-40 per image) over three representative images from each condition. The sum of the score was divided by the total number of colonies scored, and then averaged across the three images to produce a final score.

### Analysis of differentiation potential using embryoid bodies

After 6 days of differentiation in adherent culture, cells were passaged and 1.5 x 10^6^ were transferred to suspension culture plates and allowed to spontaneously differentiate without LIF or L-proline as EBs. EBs were collected at days 2, 3 and 4 and analysed with qRT-PCR for expression of the primitive streak marker *Brachyury* (*T*; primer sequences are provided in **Table S3**).

### Gene expression analysis using qRT-PCR

Total RNA was extracted from cells using GeneElute Mammalian Total RNA MiniPrep Kit (Sigma), including on-column Dnase treatment to remove any contaminating DNA. RNA was converted to cDNA using High Capacity cDNA Reverse Transcriptase Kit (Applied Biosystems). qPCR was run on 10 μL reaction volumes containing 3 μL 0.5 ng/ μL cDNA, 2 μL 1 μM primer (equal mix of Forward and Reverse primers; **Table S2**) and 5 μL 2x SYBR Green master mix (Sigma) in a 384-well plate using a Roche LightCycler 480 with the following parameters: 15 min at 95 °C, followed by 40 cycles of 30 s at 95 °C, 60 s at 60 °C, 30 s at 72 °C. Thermal melt curves were obtained following this by ramping from 60–95 °C at 2.5 °C/s. Threshold (C_t_) values were used to calculate relative expression to the reference gene, □-*actin*, employing REST v9 software. Results were normalised to untreated ESCs and transformed to log_2_ fold changes. All samples were tested to ensure that the C_t_ values for the reference gene were similar (20 ± 1 SD).

### Analysis of phosphorylation of signalling pathway intermediates

Cell samples were washed in ice-cold PBS and lysed (1 μL lysis buffer per 4 x 10^4^ cells) in the presence of protease and phosphatase inhibitors (**Table S4**). For data in **Fig. 1D,** cells were serum starved in 90% DMEM, 0.1% FBS, 0.1 mM β-Me for 4 h prior to sample collection.

Cell lysates were incubated on ice for 10 min and then centrifuged at 4°C at 12,000 rpm. The supernatant was loaded onto a 1.5 mm 12% polyacrylamide gel with a 4% stacking gel. Molecular weight markers (BioRad Precision Plus Protein Standards) were also loaded. Electrophoresis was carried out in a BioRad western blot chamber at 100 V for 2 h.

Following electrophoresis, proteins were transferred to 0.45 μm nitrocellulose membrane (BioRad) for 120 min at 100 V using a BioRad transfer system. The membrane was blocked overnight in Odyssey Blocking buffer (LiCor) at 4 °C, then washed 3 x 5 min in Tris buffered saline with Tween 20 (TBST) and then incubated with primary anti-phosphoprotein antibody overnight at 4 °C with rocking. Anti-β-tubulin antibody was used to stain for the reference protein. The membranes were then washed 3 x 5 min in TBST before 2 h incubation at room temperature in the dark with fluorescently labelled secondary antibody. Primary and secondary antibodies were diluted in Odyssey Blocking buffer with 0.1% (v/v) Tween 20. For details of antibodies and dilutions, see **Table S5**.

Membranes were imaged using an Odyssey Infrared Imaging system (LiCor), and the integrated intensity of each band was quantified with Image Studio software. Data were normalised to β-tubulin to correct for differences in loading, and then to untreated ESCs.

### Apoptosis and proliferation analysis using flow cytometry

Flow cytometry was performed on a FACS Calibur and the results quantified using FlowJo software. Apoptosis was assayed using detection of Annexin V. Live cells were centrifuged (1200 rpm x 2 min), washed in PBS and recentrifuged, and then resuspended in 100 μL Annexin V binding buffer with either (i) FITC-Annexin V conjugated antibody (1:33 dilution in TBST) and BD propidium iodide staining solution (BD Pharmingen) or (ii) PE-Annexin V conjugated antibody (1:33 dilution in TBST) and 7-AAD as per kit instructions (BD Pharmingen). Samples were analysed by flow cytometry within 30 min.

Proliferation was assayed using BrdU incorporation and processed using the FITC BrdU Flow Kit (BD Pharmingen). Briefly, BrdU was added to cells in culture at a final concentration of 10 μM and incubated for 1 h. Cells were passaged, washed in PBS, fixed in BD Cytofix/Cytoperm, and stored at −80°C in BrdU freezing buffer until required. Thawed samples were then stained according to the manufacturer’s instructions prior to flow cytometry.

### Statistical modelling and testing

Gene expression and emergent properties (cell number, proliferation, apoptosis and morphology) were modelled with (i) standard multiple linear regression (Vittinghoff *et al*., 2012), (ii) multiple linear regression with two-way interaction terms (Flanders, DerSimonian and Freedman, 1992), or (iii) Bayesian regularised neural network (BRANNGP, Burden and Winkler, 2008; Winkler and Burden, 2012). The R code used for modelling and generation of the figures is available <u>here</u>.

A correlation matrix was generated to assess broad relationships within the data (Hoyt, Imel and Chan, 2008). The *Hmisc* R package was used to generate Pearson correlations with significance levels based on rank correlation (*P* < 0.05). Parameters were ordered based on hierarchical clustering.

Before modelling, each inhibitor was encoded using a 1-hot descriptor (1 when present, 0 when absent). For modelling cell number, proliferation, and apoptosis, 1-hot variables were also used to represent each experimental day (either 2, 4 or 6). Each condition with an average of 3 replicates was used as input for modelling, with replicates averaged before modelling. As conditions containing both LY294002 and rapamycin resulted in cells not being viable after day 2, these were excluded from modelling on days 4 and 6. Data was subject to Shapiro-Wilks test, and morphology, apoptosis and proliferation data were transformed to improve normality. To ensure linearity, the residuals for each model were also measured for normality using a Shapiro-Wilks test. Model fitting parameters, including adjusted R^2^ and σ values can be found in **Table S1**. Adjusted R^2^ was used for comparability across modelling styles. Models with a higher R^2^ and lower σ values are considered to have better fit. F-tests were also calculated to compare linear models (**Table S2**), where a P-value < 0.05 indicates that the more complex models significantly improve the explanatory power of the model.

As this data contained all permutations and no predictive capacity was required, all the data was used to train models. To assess the range of responses from splitting the data, we generated 50 random 80% training/20% test models and profiled the range of responses seen in **Table S6**.

Models were generated using both the sparse linear regression method and sparse 3-layer feedforward neural network method; i.e., the Bayesian regularised neural network with a Gaussian prior (BRANNGP, Burden and Winkler, 2008). These were implemented in the specialised software package *Biomodeller*. The latter method automatically optimises the complexity of the model (number of weights) to maximise predictivity. Models were trained until the maximum of the evidence for the model so no validation set was required to provide a stopping criterion (used to denote when network training should cease), important given the small data set sizes. These models employed two neurons in the hidden layer, linear transfer functions in the input neurons and sigmoidal transfer functions in the hidden and output layer neurons. Data applied to the input layer was column scaled. See Burden and Winkler, 1999 for a detailed explanation of BRANNGP methodology.

## Supplemental Figures

**Figure S1. The P38 and PKC pathways are not active in ESCs.** Naïve ESCs were treated with either 400 μM L-proline, the P38 inhibitor SB205580 (SB, 10 μM), or staurosporine (1 mM) for the times indicated. As a positive control for P38 phosphorylation, protein lysates from the U25I human glioblastoma cell line were also used. Cell lysates were taken and analysed using western blotting. Western blots were stained for p-P38^T180/Y182^, total P38 (t-P38), p-Hsp27^S78/S82^, p-PKCζ^T410^. These samples were compared to *β*-Tubulin as a loading control.

**Figure S2. Emergent property data by inhibitor combination.** Naïve ESCs were differentiated over 6 days in 330 U/mL LIF + 400 μM L-proline with combinations of five inhibitors (U: U0126; S: SU5402; L: LY294002; R: rapamycin; P: PF-4708671). Cells were passaged at day 2, 4, and 6, and cells were counted as an indicator of cell number, and apoptosis and proliferation were measured. Morphology scoring was performed on Day 6. Conditions containing L+R (Blue) were considered non-viable at Day 2 and no data is available for this combination at Days 4 and 6. Data is shown as mean and SEM with individual data points.

**Figure S3. Correlation matrix illustrates relationships between properties.** ESCs were differentiated to EPL cells in the presence of each inhibitor combination and assayed for emergent properties (cell number, proliferation, apoptosis, morphology) and gene expression, at the days shown. **A.** Correlation matrix of all parameters (n ≥ 3), sorted by hierarchical clustering. **B.** The correlation matrix was generated using a subset of the data in panel A, along with paired data from the functional assay. The sum of the three days of *Brachyury* data was used as a proxy for the slope. Dot size and colour indicate the strength of either a positive (Blue) or negative (red) correlation.

**Figure S4. Alternative model for proliferation data.** Naïve ESCs were differentiated over 6 days in 330 U/mL LIF + 400 μM L-proline with combinations of five inhibitors (U: U0126; S: SU5402; L: LY294002; R: rapamycin; P: PF-4708671). At days 2, 4 and 6 proliferation was measured. Data was averaged across biological replicates where n ≥ 3. To correct bimodal input data, data was binned into octiles each representing an equal proportion of the data. **A.** Data was modelled using either linear modelling, linear modelling with two-way interaction terms or a Bayesian regularised neural network (BRANNGP). Fit of each model is shown comparing the actual fit with the prediction from the model. **B.** Coefficients for each variable ± SEM for standard linear model. **C.** Coefficients for each variable ± SEM for linear model with two-way interaction terms. Significance is denoted as *\*P* < 0.05.

**Figure S5. Gene expression by inhibitor combination.** Naïve ESCs were differentiated over 6 days in 330 U/mL LIF + 400 μM L-proline with combinations of five inhibitors (U: U0126; S: SU5402; L: LY294002; R: rapamycin; P: PF-4708671). At day 6, cells were collected and analysed with qRT-PCR for pluripotency genes (*Rex1* and *Oct4*, primitive ectoderm markers (*Dnmt3b, Fgf5, Lefty2* and *Otx2*), and mesendoderm *genes (Mixl1*). Data is normalized to *β*-Actin and to cells grown in 1000 U/mL LIF. No data is available for conditions containing L+R as they were considered non-viable at Day 2. Data is shown as mean log_2_ fold change and SEM with individual data points.

**Figure S6. Two-way interaction models for gene expression data.** Naïve ESCs were differentiated over 6 days in 330 U/mL LIF + 400 μM L-proline with combinations of five inhibitors (U: U0126; S: SU5402; L: LY294002; R: rapamycin; P: PF-4708671). At day 6, changes in expression of pluripotency genes (*Rex1*, **A** and *Oct4*, **B**), primitive ectoderm markers (*Dnmt3b*, **C**, *Fgf5*, **D**, *Lefty2*, **E** and *Otx2*, **F**), and mesendoderm *genes (Mixl1*, **G**). All samples were normalised to /3-Actin and then to cells grown in 1000 U/mL LIF. Data shown is for n ≥ 3 biological replicates. Data was modelled using linear modelling with two-way interaction terms. Data shows coefficients for each inhibitor ± SEM. Significance is denoted as \**P* < 0.05.

**Figure S7. Functional assay to determine position on the pluripotency continuum.** Naïve ESCs were left maintained in 1000 U/mL LIF or differentiated over 6 days in 330 U/mL LIF + 400 μM L-proline with combinations of five inhibitors (U: U0126; S: SU5402; L: LY294002; R: rapamycin; P: PF-4708671). Day 6 cells were spontaneously differentiated as embryoid bodies (EBs) on low adhesion plates in 0 U/mL LIF. mRNA samples were taken on day 2, 3 and 4 (EB2-4). Samples were analysed using qRT-PCR for *Brachyury* expression, a marker for the primitive streak. All samples were normalised to *β-Actin* as the reference gene and then to naïve ESCs. **A.** Mean log_2_ fold changes are shown ± SEM with individual data points. Data were analysed using a one-way ANOVA with *post hoc* Dunnett’s multiple comparison test to naïve ESCs. Significance is denoted as \**P* < 0.05. **Bi.** Frequency of the first significant upregulation across all inhibitor treated conditions. **Bii.** The change in frequency of first significant upregulation sorted for each condition

## Notes

### Competing Interest Statement

The authors have declared no competing interest.

## References

Acampora, D., Di Giovannantonio, L. G. and Simeone, A. (2013) ‘Otx2 is an intrinsic determinant of the embryonic stem cell state and is required for transition to a stable epiblast stem cell condition’, Development (Cambridge), 140(1), pp. 43–55. doi: 10.1242/dev.085290.

Aguilar, J. and Reyley, M. (2005) ‘The uterine tubal fluid: secretion, composition and biological effects’, Animal reproduction science, 2(2), pp. 91–105.

Aksamitiene, E., Kiyatkin, A. and Kholodenko, B. N. (2012) ‘Cross-talk between mitogenic Ras/MAPK and survival PI3K/Akt pathways: A fine balance’, Biochemical Society Transactions, 40(1), pp. 139–146. doi: 10.1042/BST20110609.

Arkun, Y. (2016) ‘Dynamic modeling and analysis of the cross-talk between insulin/akt and mapk/erk signaling pathways’, PLoS ONE, 11(3), pp. 1–22. doi: 10.1371/journal.pone.0149684.

Audet, J. (2010) ‘Adventures in time and space: Nonlinearity and complexity of cytokine effects on stem cell fate decisions’, Biotechnology and Bioengineering, 106(2), pp. 173–182. doi: 10.1002/bit.22708.

Bahrami, M., Morris, M. B. and Day, M. L. (2019) ‘Amino acid supplementation of a simple inorganic salt solution supports efficient in vitro maturation (IVM) of bovine oocytes’, Scientific Reports 2019 9:1, 9(1), pp. 1–10. doi: 10.1038/s41598-019-48038-y.

Bazer, F. W., Johnson, G. A. and Wu, G. (2015) ‘Amino acids and conceptus development during the peri-implantation period of pregnancy’, Advances in Experimental Medicine and Biology, 843, pp. 23–52. doi: 10.1007/978-1-4939-2480-6_2/COVER.

Brennan, J. et al. (2001) ‘Nodal signalling in the epiblast patterns the early mouse embryo’, Nature 2001 411:6840, 411(6840), pp. 965–969. doi: 10.1038/35082103.

Breslin, E. M. et al. (2005) ‘LY294002 and rapamycin co-operate to inhibit T-cell proliferation’, British Journal of Pharmacology, 144(6), pp. 791–800. doi: 10.1038/sj.bjp.0706061.

Burden, F. R. and Winkler, D. A. (1999) ‘Robust QSAR models using bayesian regularized neural networks’, Journal of Medicinal Chemistry, 42(16), pp. 3183–3187. doi: 10.1021/JM980697N/ASSET/IMAGES/MEDIUM/JM980697NN00001.GIF.

Burden, F. and Winkler, D. (2008) ‘Bayesian regularization of neural networks’, Methods in Molecular Biology, 458, pp. 22–44. doi: 10.1007/978-1-60327-101-1_3.

Casalino, L. et al. (2011) ‘Control of embryonic stem cell metastability by L-proline catabolism’, Journal of Molecular Cell Biology, 3(2), pp. 108–122. doi: 10.1093/jmcb/mjr001.

Cetin, I. et al. (2005) ‘Maternal and fetal amino acid concentrations in normal pregnancies and in pregnancies with gestational diabetes mellitus’, American Journal of Obstetrics and Gynecology, 192(2), pp. 610–617. doi: 10.1016/j.ajog.2004.08.011.

Chang, F. et al. (2003) ‘Involvement of PI3K/Akt pathway in cell cycle progression, apoptosis, and neoplastic transformation: A target for cancer chemotherapy’, Leukemia, 17(3), pp. 590–603. doi: 10.1038/sj.leu.2402824.

Chang, K. H. and Zandstra, P. W. (2004) ‘Quantitative screening of embryonic stem cell differentiation: Endoderm formation as a model’, Biotechnology and Bioengineering, 88(3), pp. 287–298. doi: 10.1002/bit.20242.

Chen, H. et al. (2015) ‘Erk signaling is indispensable for genomic stability and self-renewal of mouse embryonic stem cells.’, Proceedings of the National Academy of Sciences, 112(44), pp. E5936–43. doi: 10.1073/pnas.1516319112.

Cherepkova, M. Y., Sineva, G. S. and Pospelov, V. A. (2016) ‘Leukemia inhibitory factor (LIF) withdrawal activates mTOR signaling pathway in mouse embryonic stem cells through the MEK/ERK/TSC2 pathway’, Cell Death and Disease, 7, p. e2050. doi: 10.1038/cddis.2015.387.

Comes, S. et al. (2013) ‘L-proline induces a mesenchymal-like invasive program in embryonic stem cells by remodeling H3K9 and H3K36 methylation’, Stem Cell Reports, 1(4), pp. 307–321. doi: 10.1016/j.stemcr.2013.09.001.

Coucouvanis, E. and Martin, G. R. (1999) ‘BMP signaling plays a role in visceral endoderm differentiation and cavitation in the early mouse embryo’, Development (Cambridge, England), 126(3), pp. 535–546. doi: 10.1242/DEV.126.3.535.

D’Aniello, C. et al. (2015) ‘A novel autoregulatory loop between the Gcn2-Atf4 pathway and L-Proline metabolism controls stem cell identity’, Cell Death and Differentiation, 22(7), pp. 1094–1105. doi: 10.1038/cdd.2015.24.

D’Aniello, C. et al. (2017) ‘Vitamin C and L-Proline antagonistic effects capture alternative states in the pluripotency continuum’, Stem Cell Reports, 8(1), pp. 1–10. doi: 10.1016/j.stemcr.2016.11.011.

Dailey, L. et al. (2005) ‘Mechanisms underlying differential responses to FGF signaling’, Cytokine and Growth Factor Reviews, 16(2 SPEC. ISS.), pp. 233–247. doi: 10.1016/j.cytogfr.2005.01.007.

Doetschman, T. C. et al. (1985) ‘The *in vitro* development of blastocyst-derived embryonic stem cell lines: formation of visceral yolk sac, blood islands and myocardium’, Journal of Embryology and Experimental Morphology, 87(1), pp. 27–45.

Dowling, R. J. O. et al. (2010) ‘mTORC1-mediated cell proliferation, but not cell growth, controlled by the 4E-BPs’, Science (New York, N.Y.), 328(5982), p. 1172. doi: 10.1126/SCIENCE.1187532.

Epa, V. C. et al. (2013) ‘Europe PMC Funders Group Modelling human embryoid body cell adhesion to a combinatorial library of polymer surfaces’, 22(39), pp. 20902–20906. doi: 10.1039/C2JM34782B.Modelling.

Favata, M. F. et al. (1998) ‘Identification of a novel inhibitor of mitogen-activated protein kinase kinase’, Journal of Biological Chemistry, 273(29), pp. 18623–18632. doi: 10.1074/jbc.273.29.18623.

Fingar, D. C. et al. (2002) ‘Mammalian cell size is controlled by mTOR and its downstream targets S6K1 and 4EBP1/eIF4E.’, Genes and Development, 16(12), pp. 1472–87. doi: 10.1101/gad.995802.

Flanders, W. D., DerSimonian, R. and Freedman, D. S. (1992) ‘Interpretation of linear regression models that include transformations or interaction terms’, Annals of Epidemiology, 2(5), pp. 735–744. doi: 10.1016/1047-2797(92)90018-L.

Gardner, D. K. and Lane, M. (1993) ‘Amino acids and ammonium regulate mouse embryo development in culture.’, Biology of Reproduction, 48(2), pp. 377–85.

Glover, H. J., Shparberg, R. A. and Morris, M. B. (2022) ‘L-Proline Supplementation Drives Self-Renewing Mouse Embryonic Stem Cells to a Partially Primed Pluripotent State: The Early Primitive Ectoderm-Like Cell’, Methods in molecular biology (Clifton, N.J.), 2490, pp. 11–24. doi: 10.1007/978-1-0716-2281-0_2.

Gross, V. S., Hess, M. and Cooper, G. M. (2005) ‘Mouse embryonic stem cells and preimplantation embryos require signaling through the phosphatidylinositol 3-kinase pathway to suppress apoptosis’, Molecular Reproduction and Development, 70(3), pp. 324–332. doi: 10.1002/mrd.20212.

Harris, S. E. et al. (2005) ‘Nutrient concentrations in murine follicular fluid and the female reproductive tract.’, Theriogenology, 64(4), pp. 992–1006. doi: 10.1016/j.theriogenology.2005.01.004.

Hart, A. H. et al. (2002) ‘Mixl1 is required for axial mesendoderm morphogenesis and patterning in the murine embryo’, Development (Cambridge, England), 129(15), pp. 3597–3608. doi: 10.1242/DEV.129.15.3597.

Harvey, N. T. et al. (2010) ‘Response to BMP4 signalling during ES cell differentiation defines intermediates of the ectoderm lineage.’, Journal of Cell Science, 123(Pt 10), pp. 1796–804. doi: 10.1242/jcs.047530.

Hoogland, S. H. A. and Marks, H. (2021) ‘Developments in pluripotency: a new formative state’, Cell research, 31(5), pp. 493–494. doi: 10.1038/S41422-021-00494-W.

Hoyt, W. T., Imel, Z. E. and Chan, F. (2008) ‘Multiple Regression and Correlation Techniques: Recent Controversies and Best Practices’, Rehabilitation Psychology, 53(3), pp. 321–339. doi: 10.1037/A0013021.

Huang, G. et al. (2014) ‘STAT3 Phosphorylation at Tyrosine 705 and Serine 727 Differentially Regulates Mouse ESC Fates’, Stem cells (Dayton, Ohio), 32(5), p. 1149. doi: 10.1002/STEM.1609.

Ireland, R. G. et al. (2020) ‘Combinatorial extracellular matrix microarray identifies novel bioengineered substrates for xeno-free culture of human pluripotent stem cells’, Biomaterials, 248(August 2019), p. 120017. doi: 10.1016/j.biomaterials.2020.120017.

Jakobsen, R. B. et al. (2014) ‘Analysis of the effects of five factors relevant to in vitro chondrogenesis of human mesenchymal stem cells using factorial design and high throughput mRNA-profiling’, PLoS ONE, 9(5). doi: 10.1371/journal.pone.0096615.

Jirmanova, L. et al. (2002) ‘Differential contributions of ERK and PI3-kinase to the regulation of cyclin D1 expression and to the control of the G1/S transition in mouse embryonic stem cells.’, Oncogene, 21(36), pp. 5515–5528. doi: 10.1038/sj.onc.1205728.

Julkunen, H. et al. (2020) ‘Leveraging multi-way interactions for systematic prediction of pre-clinical drug combination effects’, Nature Communications, 11(1). doi: 10.1038/s41467-020-19950-z.

Kim, J. et al. (2008) ‘An Extended Transcriptional Network for Pluripotency of Embryonic Stem Cells’, Cell, 132(6), pp. 1049–1061. doi: 10.1016/j.cell.2008.02.039.

Kunath, T. et al. (2007) ‘FGF stimulation of the Erk1/2 signalling cascade triggers transition of pluripotent embryonic stem cells from self-renewal to lineage commitment.’, Development, 134(16), pp. 2895–902. doi: 10.1242/dev.02880.

Lane, M. and Gardner, D. K. (1997) ‘Nonessential amino acids and glutamine decrease the time of the first three cleavage divisions and increase compaction of mouse zygotes in vitro.’, Journal of Assisted Reproduction and Genetics, 14(7), pp. 398–403. doi: 10.1007/BF02766148.

Lanner, F. and Rossant, J. (2010) ‘The role of FGF/Erk signaling in pluripotent cells’, Development, 137(20), pp. 3351–3360. doi: 10.1242/dev.050146.

Lonic, A. (2006) Molecular mechanism of L-proline induced EPL-cell formation. University of Adelaide.

Mendoza, M. C., Er, E. E. and Blenis, J. (2011) ‘The Ras-ERK and PI3K-mTOR pathways: Cross-talk and compensation’, Trends in Biochemical Sciences, 36(6), pp. 320–328. doi: 10.1016/j.tibs.2011.03.006.

Minchiotti, G. et al. (2022) ‘Capturing Transitional Pluripotency through Proline Metabolism’, Cells, 11(14). doi: 10.3390/CELLS11142125.

Mohammadi, M. et al. (1997) ‘Structures of the tyrosine kinase domain of fibroblast growth factor receptor in complex with inhibitors’, Science, 276(5314), pp. 955–960. doi: 10.1126/SCIENCE.276.5314.955/ASSET/C77197A2-82FE-4E7E-A3C4-E4212C555522/ASSETS/GRAPHIC/SE1975130007.JPEG.

Morgani, S., Nichols, J. and Hadjantonakis, A. K. (2017) ‘The many faces of Pluripotency: In vitro adaptations of a continuum of in vivo states’, BMC Developmental Biology, 17(1), pp. 10–12. doi: 10.1186/s12861-017-0150-4.

Morris, M. B. et al. (2020) ‘Selected Amino Acids Promote Mouse Pre-implantation Embryo Development in a Growth Factor-Like Manner’, Frontiers in Physiology, 11(March), pp. 1–12. doi: 10.3389/fphys.2020.00140.

Murakami, M. et al. (2004) ‘mTOR is essential for growth and proliferation in early mouse embryos and embryonic stem cells.’, Molecular and Cellular Biology, 24(15), pp. 6710–8. doi: 10.1128/MCB.24.15.6710-6718.2004.

Nawroth, R. et al. (2011) ‘S6K1 and 4E-BP1 Are Independent Regulated and Control Cellular Growth in Bladder Cancer’, PLOS ONE, 6(11), p. e27509. doi: 10.1371/JOURNAL.PONE.0027509.

Panina, S. B. et al. (2020) ‘Utilizing Synergistic Potential of Mitochondria-Targeting Drugs for Leukemia Therapy’, Frontiers in Oncology, 10(April). doi: 10.3389/fonc.2020.00435.

Patriarca, E. J. et al. (2021) ‘The Multifaceted Roles of Proline in Cell Behavior’, Frontiers in cell and developmental biology, 9. doi: 10.3389/FCELL.2021.728576.

Pearce, L. R. et al. (2010) ‘Characterization of PF-4708671, a novel and highly specific inhibitor of p70 ribosomal S6 kinase (S6K1).’, The Biochemical Journal, 431(2), pp. 245–55. doi: 10.1042/BJ20101024.

Pelton, T. A. et al. (2002) ‘Transient pluripotent cell populations during primitive ectoderm formation: correlation of *in vivo* and *in vitro* pluripotent cell development.’, Journal of Cell Science, 115(Pt 2), pp. 329–339.

Pons, B. et al. (2011) ‘The effect of p-4E-BP1 and p-eIF4E on cell proliferation in a breast cancer model’, International journal of oncology, 39(5), pp. 1337–1345. doi: 10.3892/IJO.2011.1118.

Prudhomme, W. A., Duggar, K. H. and Lauffenburger, D. A. (2004) ‘Cell population dynamics model for deconvolution of murine embryonic stem cell self-renewal and differentiation responses to cytokines and extracellular matrix’, Biotechnology and Bioengineering, 88(3), pp. 264–272. doi: 10.1002/bit.20244.

Rathjen, J. et al. (1999) ‘Formation of a primitive ectoderm like cell population, EPL cells, from ES cells in response to biologically derived factors.’, Journal of Cell Science, 112(5), pp. 601–612.

Rathjen, J. et al. (2002) ‘Directed differentiation of pluripotent cells to neural lineages: Homogeneous formation and differentiation of a neurectoderm population’, Development,129(11), pp. 2649–2661. doi: 10.1242/dev.129.11.2649.

Rivera-Pérez, J. A. and Hadjantonakis, A. K. (2015) ‘The Dynamics of Morphogenesis in the Early Mouse Embryo’, Cold Spring Harbor Perspectives in Biology, 7(11). doi: 10.1101/CSHPERSPECT.A015867.

Sabers, C. J. et al. (1995) ‘Isolation of a protein target of the FKBP12-rapamycin complex in mammalian cells’, The Journal of biological chemistry, 270(2), pp. 815–822. doi: 10.1074/JBC.270.2.815.

Shparberg, R. A., Glover, H. J. and Morris, M. B. (2019) ‘Embryoid body differentiation of mouse embryonic stem cells into neurectoderm and neural progenitors’, Methods in Molecular Biology, 2029, pp. 273–285. doi: 10.1007/978-1-4939-9631-5_21.

Shparberg, R., Glover, H. and Morris, M. B. (2019) ‘Modeling Mammalian Commitment to the Neural Lineage Using Embryos and Embryonic Stem Cells’, Frontiers in physiology, 10(MAY). doi: 10.3389/FPHYS.2019.00705.

Smith, A. (2017) ‘Formative pluripotency: the executive phase in a developmental continuum.’, Development, 144(3), pp. 365–373. doi: 10.1242/dev.142679.

Snow, M. (1977) ‘Gastrulation in the mouse: Growth and regionalization of the epiblast’, J. Embryol. exp. Morph, 42, pp. 293–303.

Sorokin, M. et al. (2018) ‘Oncobox bioinformatical platform for selecting potentially effective combinations of target cancer drugs using high-throughput gene expression data’, Cancers, 10(10). doi: 10.3390/cancers10100365.

Stavridis, M. P., Collins, B. J. and Storey, K. G. (2010) ‘Retinoic acid orchestrates fibroblast growth factor signalling to drive embryonic stem cell differentiation’, Development, 137(6), pp. 881–890. doi: 10.1242/dev.043117.

Stead, E. et al. (2002) ‘Pluripotent cell division cycles are driven by ectopic Cdk2, cyclin A/E and E2F activities.’, Oncogene, 21(54), pp. 8320–33. doi: 10.1038/sj.onc.1206015.

Takahashi, K., Murakami, M. and Yamanaka, S. (2005) ‘Role of the phosphoinositide 3-kinase pathway in mouse embryonic stem (ES) cells’, Biochemical Society Transactions, 33(6), pp. 1522–1525. doi: 10.1042/BST20051522.

Takeuchi, H. et al. (2005) ‘Synergistic augmentation of rapamycin-induced autophagy in malignant glioma cells by phosphatidylinositol 3-kinase/protein kinase B inhibitors’, Cancer Research, 65(8), pp. 3336–3346. doi: 10.1158/0008-5472.CAN-04-3640.

Tan, B. S. N. et al. (2011) ‘The amino acid transporter SNAT2 mediates l-proline-induced differentiation of ES cells’, American Journal of Physiology - Cell Physiology, 300(6). doi: 10.1152/ajpcell.00235.2010.

Tan, B. S. N. et al. (2016) ‘Src family kinases and p38 Mitogen-Activated Protein Kinases regulate pluripotent cell differentiation in culture’, PLoS One. Edited by M. Schubert, 11(10), p. e0163244. doi: 10.1371/journal.pone.0163244.

Treleaven, T. et al. (2021) ‘In vitro fertilisation of mouse oocytes in l-proline and l-pipecolic acid improves subsequent development’, Cells, 10(6). doi: 10.3390/cells10061352.

Tsurutani, J. et al. (2005) ‘Inhibition of the phosphatidylinositol 3-kinase/Akt/mammalian target of rapamycin pathway but not the MEK/ERK pathway attenuates laminin-mediated small cell lung cancer cellular survival and resistance to imatinib mesylate or chemotherapy’, Cancer Research, 65(18), pp. 8423–8432. doi: 10.1158/0008-5472.CAN-05-0058.

Vittinghoff, E. et al. (2012) ‘Regression Methods in Biostatistics’. doi: 10.1007/978-1-4614-1353-0.

Vlahos, C. J. et al. (1994) ‘A specific inhibitor of phosphatidylinositol 3-kinase, 2-(4-morpholinyl)- 8-phenyl-4H-1-benzopyran-4-one (LY294002)’, Journal of Biological Chemistry, 269(7), pp. 5241–5248. doi: 10.1016/s0021-9258(17)37680-9.

Wang, C. et al. (2013) ‘Functional crosstalk between AKT/mTOR and Ras/MAPK pathways in hepatocarcinogenesis: Implications for the treatment of human liver cancer’, Cell Cycle, 12(13), pp. 1999–2010. doi: 10.4161/cc.25099.

Wang, X et al. (2021) ‘Formative pluripotent stem cells show features of epiblast cells poised for gastrulation’, Cell research, 31(5), pp. 526–541. doi: 10.1038/S41422-021-00477-X.

Washington, J. M. et al. (2010) ‘L-Proline induces differentiation of ES cells: a novel role for an amino acid in the regulation of pluripotent cells in culture’, American Journal of Physiology - Cell Physiology, 298, pp. 982–992.

Watanabe, D. et al. (2002) ‘Stage- and cell-specific expression of Dnmt3a and Dnmt3b during embryogenesis’, Mechanisms of Development, 118(1-2), pp. 187–190. doi: 10.1016/S0925-4773(02)00242-3.

Van Winkle, L. J. (2001) ‘Amino Acid Transport Regulation and Early Embryo Development’, Biology of Reproduction, 64(1), pp. 1–12. doi: 10.1095/BIOLREPROD64.1.1.

Van Winkle, L. J. et al. (2006) ‘System B0,+ amino acid transport regulates the penetration stage of blastocyst implantation with possible long-term developmental consequences through adulthood’, Human reproduction update, 12(2), pp. 145–157. doi: 10.1093/HUMUPD/DMI044.

Winkler, D. A. and Burden, F. R. (2012) ‘Robust, quantitative tools for modelling ex-vivo expansion of haematopoietic stem cells and progenitors’, Molecular BioSystems, 8(3), pp. 913–920. doi: 10.1039/c2mb05439f.

Woolf, P. J. et al. (2005) ‘Bayesian analysis of signaling networks governing embryonic stem cell fate decisions’, Bioinformatics, 21(6), pp. 741–753. doi: 10.1093/bioinformatics/bti056.

Yellen, P. et al. (2011) ‘High-dose rapamycin induces apoptosis in human cancer cells by dissociating mTOR complex 1 and suppressing phosphorylation of 4E-BP1’, http://dx.doi.org/10.4161/cc.10.22.18124, 10(22), pp. 3948–3956. doi: 10.4161/CC.10.22.18124.

Yu, J. S. L. and Cui, W. (2016) ‘Proliferation, survival and metabolism: The role of PI3K/AKT/ mTOR signalling in pluripotency and cell fate determination’, Development (Cambridge), 143(17), pp. 3050–3060. doi: 10.1242/dev.137075.

